## Supplementary material for "Fibrous Network Nature of Plant Cell Walls Enables Tunable Mechanics for Development": SI

### Mechanical properties are tuned during development with the fibrous network nature of the Arabidopsis cell wall

#### Supporting Information

##### A Two-dimensional affine isotropic network model

The two dimensional affine isotropic network model [1] is based on averaging the response of a single fiber to an applied deformation over all orientations of randomly distributed fibers in a plane. We assume that all fibers only support force along their current orientation axis (i.e. no shear) and are only able to stretch or compress in that same axis. A fiber with initial orientation $\hat{n} = (\cos\theta, \sin\theta)$, which is subjected to deformation gradient tensor $\mathbf{\Lambda}$, will deform axially and rotate. The force within the fiber is given by $k(|\mathbf{\Lambda}\hat{\mathbf{n}}| - 1)$, where $k$ is the stiffness of the fiber and set to 1 for simplicity. The new orientation of the fiber would be $(\mathbf{\Lambda}\hat{\mathbf{n}})/|\mathbf{\Lambda}\hat{\mathbf{n}}|$. Taking the average over all orientations, we have the membrane stress tensor of the network:

$$\begin{aligned}\sigma_{ij} &= \frac{\rho}{\det \mathbf{\Lambda}} \left\langle k(|\mathbf{\Lambda}\hat{\mathbf{n}}| - 1)\frac{\Lambda_{il} n_l \Lambda_{jm} n_m}{|\mathbf{\Lambda}\hat{\mathbf{n}}|} \right\rangle \\ &= \frac{\rho}{2\pi \det \mathbf{\Lambda}} \int_\theta k(|\mathbf{\Lambda}\hat{\mathbf{n}}| - 1)\frac{\Lambda_{il} n_l \Lambda_{jm} n_m}{|\mathbf{\Lambda}\hat{\mathbf{n}}|} d\theta \end{aligned} \tag{1}$$

where $\rho$ is fiber length density (length per unit of area), set to $\frac{1}{2\pi}$ for simplicity.

In the uniaxial tension case, consider applying $\mathbf{\Lambda} = \begin{pmatrix} \lambda_1 & 0 \\ 0 & \lambda_2 \end{pmatrix}$, where $\lambda_1$ is prescribed value and $\lambda_2$ needs to be solved such that $\sigma_{22} = 0$. Here

$$|\mathbf{\Lambda}\hat{\mathbf{n}}| = \sqrt{\lambda_1^2 \cos^2\theta + \lambda_2^2 \sin^2\theta} \tag{2}$$

$$\det \mathbf{\Lambda} = \lambda_1 \lambda_2 \tag{3}$$

The stress tensor for the fiber network is then given by:

$$\sigma_{11} = \frac{1}{\det \mathbf{\Lambda}} \int_\theta \frac{(|\mathbf{\Lambda}\hat{\mathbf{n}}| - 1)\lambda_1^2 \cos^2\theta}{|\mathbf{\Lambda}\hat{\mathbf{n}}|} d\theta \tag{4}$$

$$\sigma_{22} = \frac{1}{\det \mathbf{\Lambda}} \int_\theta \frac{(|\mathbf{\Lambda}\hat{\mathbf{n}}| - 1)\lambda_2^2 \sin^2\theta}{|\mathbf{\Lambda}\hat{\mathbf{n}}|} d\theta \tag{5}$$

$$\sigma_{12} = \frac{1}{\det \mathbf{\Lambda}} \int_\theta \frac{(|\mathbf{\Lambda}\hat{\mathbf{n}}| - 1)\lambda_1 \lambda_2 \cos\theta \sin\theta}{|\mathbf{\Lambda}\hat{\mathbf{n}}|} d\theta \tag{6}$$

To solve $\sigma_{22} = 0$ numerically, we assume two initial values for $\lambda_2$: $\lambda_{2_{t0}}$ and $\lambda_{2_{t1}}$, and calculate corresponding $\sigma_{22_{t0}}$ and $\sigma_{22_{t1}}$. We use the Secant method to find the next guess value:

$$\lambda_{2_{t2}} = \lambda_{2_{t1}} - \sigma_{22_{t1}} \frac{\lambda_{2_{t1}} - \lambda_{2_{t0}}}{\sigma_{22_{t1}} - \sigma_{22_{t0}}} \tag{7}$$

We repeat this process until $\sigma_{22} < 10^{-10}$, so that we find the $\lambda_2$. Then, the stress tensor can be calculated based on $\lambda_1$ and $\lambda_2$ using Eqns 4, 5 and 6. The membrane force (nominal stress) can be calculated by $S_{11} = \sigma_{11}\lambda_2$.

#### B Five-beam model

In the five-beam model [4], there are five beams, and they are linked by connectors that transmit force, but not moments, between beams. When the structure deforms, the four tilted beams AC, AD, BC, and BD will rotate and stretch, the beam AB will deform only through bending, and the connectors will undergo shear deformation and slide along the beam if the load exceeds a certain threshold (Figure 8A). We first look at the force-deformation relationship of each component respectively.

###### Stretching beam with stretching stiffness $k_s$

These beams are elongated elastically under stretching force $F_s$. The elongation length is given by $\Delta L_s = F_s/k_s$ (Figure 8B).

###### Bending Beam with bending stiffness $k_b$

This beam is bent elastically under compression force $F_b$. The end of beam moves distance $\Delta L_b = F_b/k_b$(Figure 8C).

###### Connector with shear stiffness $k_c$ and sliding resistance $D$

The connector is under shear when two connected beams are pulled apart from each other. Two kinds of displacement need to be considered in this case. First, there is elastic deformation of the connector resulting in displacement $\Delta u_c = F_c/k_c$. Then, fibers are allowed to slide relative to each other along the connector, resulting in displacement $\Delta u_{slide} = (F_c - F_0)/D$, only when the force $F_c$ exceeds the threshold $F_0$. During unloading the elastic deformation of the connector can fully recover, while the sliding distance cannot recover (*i.e.,* $\Delta u$ is set according to the maximum $F_c$ value experience during the loading history). Therefore, the total displacement is (Figure 8D): If reloading, the sliding distance remains unchanged until the load exceeds the maximum load $F_m$ in history. The maximum load $F_m$ serves as a new yield force threshold. After exceeding the maximum load, the sliding distance still follows the equation: $\Delta u_{slide} = (F_c - F_0)/D$. Therefore, the total displacement

$$\Delta u_c + \Delta u_{slide} = h(F_c) = \begin{cases} \frac{F_c}{k_c} + \text{const} & \text{if during unloading or if loading with } F_c < F_m \\ F_c(\frac{1}{k_c} + \frac{1}{D}) - \frac{F_0}{D} & \text{if loading with } F_c > F_m \end{cases} \tag{8}$$

where $h$ is the piecewise function.
Consider a five-beam structure, the initial length of each of the four bending beams is the same, denoted as $L_{b0}$, and the initial tilt angle of the stretching beam is $\theta_0$, so the initial length of each the stretching beams is $L_{s0} = \frac{L_{b0}}{2\sin\theta_0}$. A force $F$ in x direction is prescribed on the node C and D of the five-beam structure. According to force balance relations, the same force acts on each of the stretching beams:

$$F_s = \frac{F}{2\cos\theta} \tag{9}$$

and the force acting on the bending beam:

$$F_b = F\tan\theta \tag{10}$$

and the force acting on the connectors:

$$F_c = F_s = \frac{F}{2\cos\theta} \tag{11}$$

where $\theta$ is the current tilt angle of the stretching beams.
According to the geometric relation, we have the distance between nodes A and C:

$$L_{AC} = L_{s0} + \Delta L_s + \Delta u_c + \Delta u_{slide} \tag{12}$$

the distance between nodes A and B:

$$L_{AB} = L_{b0} - \Delta L_b - \Delta u_c - \Delta u_{slide} \tag{13}$$

the distance between node C and D:

$$L_{CD} = 2L_{AC}\cos\theta \tag{14}$$

$L_{AB}$ and $L_{AC}$ should satisfy

$$2L_{AC}\sin\theta = L_{AB} \tag{15}$$

According to force-displacement relations discussed above, we have

$$\Delta L_s = F_s/k_s \tag{16}$$

$$\Delta L_b = F_b/k_b \tag{17}$$

$$\Delta u_c + \Delta u_{slide} = h(F_c) \tag{18}$$

So far we have 10 unknowns: $\theta$, $F_s$, $F_b$, $F_c$, $\Delta L_s$, $\Delta L_b$, $\Delta u_c + \Delta u_{slide}$, $L_{AC}$, $L_{AB}$, $L_{CD}$, which can be solved from the above ten Eqns 9-18.
To increment the applied force $F$ by $dF$ in each step, we use the incremental form of these equations:

$$dF = 2\cos\theta dF_s - 2F_s \sin\theta d\theta \tag{19}$$

$$dF_b = dF\tan\theta + \frac{Fd\theta}{\cos^2\theta} \tag{20}$$

$$dF_c = dF_s \tag{21}$$

$$dL_{AC} = d\Delta L_s + d\Delta u_c + d\Delta u_{slide} \tag{22}$$

$$dL_{AB} = -d\Delta L_b - d\Delta u_c - d\Delta u_{slide} \tag{23}$$

$$dL_{CD} = 2dL_{AC}\cos\theta - 2L_{AC}\sin\theta d\theta \tag{24}$$

$$2dL_{AC}\sin\theta + 2L_{AC}\cos\theta d\theta = dL_{AB} \tag{25}$$

$$d\Delta L_s = \frac{dF_s}{k_s} \tag{26}$$

$$d\Delta L_b = \frac{dF_b}{k_b} \tag{27}$$

$$d\Delta u_c + d\Delta u_{slide} = \frac{dh(F_c)}{dF_c} dF_c \tag{28}$$

Combining Eqns 19 - 28, we obtain:

$$-\frac{dF_b}{k_b} - \frac{dh(F_b)}{dF_b} dF_b = 2\left(\frac{dF_s}{k_s} + \frac{dh(F_s)}{dF_s} dF_s\right)\sin\theta + 2L_{AB}\cos\theta d\theta \tag{29}$$

Note here $\theta$ and $L_{AC}$ are measured in current configuration after $dF$ is loaded in current step, and can be estimated as $\theta_p$ and $L_{AC_p}$ in the previous step if $dF$ is very small. Then $dF_b$, $dF_s$ and $d\theta$ can be solved from Eqns 19, 21 and 29.
Finally, the incremental stretches of the five-beam structure can be calculated as:

$$d\lambda_x = \frac{dL_{CD}}{L_{CD_0}} = \frac{dL_{AC}\cos\theta - L_{AC}\sin\theta d\theta}{L_{s0}\cos\theta_0} \tag{30}$$

$$d\lambda_y = \frac{dL_{AB}}{L_{b0}} \tag{31}$$

with total stretches of:

$$\lambda_x = \frac{L_{AC}\cos\theta}{L_{s0}\cos\theta_0} - 1 \tag{32}$$

$$\lambda_y = \frac{L_{AB}}{L_{b0}} - 1 \tag{33}$$

In plant cell walls, the distance between adjacent cellulose microfibrils is around $100 \pm 40$nm, and the thickness of microfibrils is around $3.6 \pm 1.9$nm [2], so we chose the stretching stiffness and bending stiffness ratio $\frac{k_s}{k_b} = \frac{Al^2}{3I} \sim \frac{l^2}{d^2} \sim 100$, where $A$ and $I$ represent the area and the second moment of the area. For a qualitative comparison, we chose the following parameters. We set $k_b = 1$ so that $k_s = 100$. The length of the beam simply scales the overall force-displacement response, so we made the arbitrary choice of $l_b0 = 10$. The initial angle $\theta_0$ will influence the incremental Poisson's ratio. We chose $\theta_0 = \frac{1.3\pi}{4}$ so that the initial incremental Poisson's ratio of the model was within the same range as the experiments. We chose $k_c = 36$ and $D = 24$ so that the stiffening ratio and recovery ratio were comparable to the experiments on the 25d samples. The connector resistance was calculated as $C = \frac{1}{\frac{1}{D}+\frac{1}{k_c}}$. Different connector resistance values $C_1$, $C_2$ and $C_3$ were corresponding to three sets of chose value(1) $k_c = 15$ and $D = 10$; (2)$k_c = 30$ and $D = 20$; (3) $k_c = 36$ and $D = 24$ respectively. Alternative initial angle values of $\theta_1 = \frac{1.2\pi}{4}$ and $\theta_2 = \frac{1.4\pi}{4}$ were chosen to demonstrate possible anisotropic effects.

### C Turgor pressure and cell deformation relation under linear and stiffening behavior

Considering a spherical cell with initial radius $r_0$, assume the cell wall is isotropic, so that the cell will expand to new radius $r$ under turgor pressure $P$ (Figure 9A). The stretch ratio of the cell wall $\lambda = \frac{r}{r_0}$. According to Laplace's law, the membrane force within cell wall is

$$T = \frac{Pr}{2} \tag{34}$$

##### Case I

In linear mechanical behavior case, the membrane force within the cell wall would always be proportional to the stretch, in other word, the stiffness $E$ is constant. Therefore, we have membrane force and stretch relation as:

$$T = E(\lambda - 1) \tag{35}$$

Combining Eqns 34 and 35, we have turgor pressure and stretch relation as:

$$P = E\frac{\lambda - 1}{\lambda}\frac{2}{r_0} \tag{36}$$

##### Case II

In the nonlinear stiffening case, we consider the stiffness $E = E(\lambda)$ as a function of the stretch. Therefore, we have membrane force and stretch relation in incremental form as:

$$\delta T = E(\lambda)\delta\lambda \tag{37}$$

Combining Eqns 34 and 37, we have turgor pressure and stretch relation as incremental form as:

$$\delta P = \frac{2\delta T}{\lambda r_0} - \frac{2T\delta\lambda}{\lambda^2 r_0} \tag{38}$$

We consider the initial cell size is 10 μm. For **case I**, the stiffness $E = 14.43$(Figure 9B). For **case II**, we use a sigmoid function from fitting the stiffness versus stretch curve in Figure 1E, which is $E(\lambda) = E_0 + \frac{E_1 - E_0}{1+e^{-a(x-b)}} = 14.43 + \frac{51.71}{1+e^{-58.29(x-1.119)}}$(Figure 9B). The turgor pressure and stretch curves for both cases are plotted in Figure 9C.

#### D Mechanical behavior under general loading conditions

In this section, we will derive the deformation of the cell wall under general loading conditions using results from tensile tests. We will consider both linear and nonlinear elasticity formulations.

##### D.1 Linear isotropic elastic formulation

Linear elasticity formulation can be used under the infinitesimal deformation assumption [5], where the deformation changes linearly with applied force. Since cell wall is very thin and its mechanical properties are similar within the plane from our tensile test results, we consider two-dimensional isotropic elastic theory using plane stress state [5], the deformation under loading can be characterized from Hooke's law:

$$
\begin{aligned}
\epsilon_x &= \frac{1}{E}(s_x - \nu s_y) \\
\epsilon_y &= \frac{1}{E}(s_y - \nu s_x)
\end{aligned}
\tag{39}
$$

where $\epsilon_x$ and $\epsilon_y$ are strains in x and y directions respectively, defined as the stretch in each direction minus 1. $s_x$ and $s_y$ are membrane forces in x and y directions respectively. $E$ and $\nu$ are Young's modulus and Poisson's ratio respectively.

From tensile tests described in the main text, we have $s_y = 0$. Therefore, $E = s_x/\epsilon_x$ and $\nu = -\frac{\epsilon_y}{\epsilon_x}$. Once $E$ and $\nu$ are determined, the deformation of the cell wall ($\epsilon_x$ and $\epsilon_y$) can be calculated under any loading conditions (varying $s_x$ and $s_y$). We will consider two types of loading conditions that are closely related to what cell walls experience in a live cell and demonstrate how to calculate the resulting deformation.

**Case I: equal-biaxial loading** The cell wall of a spherical cell (radius $r$) is subjected to equal-biaxial loading due to the turgor pressure $P$. According to Laplace's law, membrane forces within the cell wall are: $s_x = s_y = \frac{Pr}{2}$. Therefore, the deformation can be calculated according to Eq. 39: $\epsilon_x = \epsilon_y = \frac{1-\nu}{E}\frac{Pr}{2}$. From this result, we can see that a larger Poisson's ratio makes cell wall more resistant to deform under equal-biaxial loading.

**Case II: turgid cylindrical cell under stretch** Considering a cylindrical cell with turgor pressure $P$, length $l_0$ and radius $r_0$. The cell is stretched by an axial force $\delta F$ (Figure 10A), resulting in length change $\delta l$ and radius change $\delta r$. Here we assume turgor pressure is kept as constant. The membrane force in axial direction $\delta T_a = \frac{P\delta r}{2} + \frac{\delta F}{2\pi r_0}$, and in transverse(circumferential) direction $\delta T_h = P\delta r$. The strain in axial direction $\epsilon_a = \frac{\delta l}{l_0}$, and in transverse direction $\epsilon_h = \frac{\delta r}{r_0}$. Combining Eq. 39, we can calculate the stiffness of this cell as: $E_{cell} = \frac{\delta F/\pi r_o^2}{\epsilon_a} = \frac{2E(2E-Pr_0(2-\nu))}{r_0(2E-Pr_0(1-\nu^2))}$. If we take Young's modulus $E = 20$ N/m, we have turgor pressure $P = 0.5$ MPa, and the radius $r = 10$ µm, and plot $E_{cell}$ versus $\nu$ (Figure 10B). From this plot, a larger Poisson's ratio leads a larger stiffness $E_{cell}$, indicating that the cell becomes more resistant to forces from its environment or neighboring cells.

##### D.2 nonlinear elastic formulation

The cell wall has a polylamellate structure [7], which results in different mechanical properties along the thickness direction compared to within the plane of the cell wall. From our tensile test results, mechanical properties are similar within the cell wall plane. Therefore, we consider a nonlinear transversely isotropic (plane isotropy) formulation based on hyperelastic constitutive theories [3,6]. Given that the cell wall matrix, composed of pectin and hemicellulose, is hydrogel-like, we assume the cell wall is incompressible (volume-conserving). We assume the strain energy(Helmholtz free energy) form [6]: $W = w(I_1, \lambda_3) - p(J-1)$, where $I_1 = \lambda_1^2 + \lambda_2^2 + \lambda_3^2$, $J = \lambda_1\lambda_2\lambda_3 = 1$, $p$ introduced serves as an indeterminate Lagrange multiplier, $\lambda_1$ and $\lambda_2$ are the stretches in two perpendicular directions within the cell wall plane, and $\lambda_3$ is the stretch along the thickness direction. The strain energy function fully characterizes the mechanical behavior, allowing us to determine the relationship between force and deformation. Therefore, we have

$$
\begin{aligned}
s_1 &= 2\lambda_1\beta - p\lambda_2\lambda_3 \\
s_2 &= 2\lambda_2\beta - p\lambda_1\lambda_3 \\
s_3 &= 2\lambda_3\beta + \alpha - p\lambda_2\lambda_3
\end{aligned}
\tag{40}
$$

where $\beta = \frac{\partial w}{\partial I_1}$ and $\alpha = \frac{\partial w}{\lambda_3}$. From tensile tests described in the main text, we have $s_2 = 0$, $s_3 = 0$. $s_1$, $\lambda_1$, and $\lambda_2$ are measured, and $\lambda_3 = \frac{1}{\lambda_1\lambda_2}$. So that we have $\beta = \frac{s_1\lambda_1}{2(\lambda_1^2-\lambda_2^2)}$ and $\alpha = \frac{2\beta(\lambda_2^2-\lambda_3^2)}{\lambda_3}$. We assume that the strain energy does not include cross-terms involving $I_1$ and $\lambda_3$, so $\beta$ only depends on $I_1$ and $\alpha$ only depends on $\lambda_3$. Therefore, these functions can be determined respectively. Then we will explore relation between force and deformation under equal-biaxial loading based on Eq. 40.

**equal-biaxial loading** We have $\lambda_1 = \lambda_2 = \lambda_b$ and $\lambda_3 = \lambda_t$. Membrane forces in plane $s_1 = s_2 = s = 2\beta_1\lambda_b - p\lambda_b\lambda_t$, where $p = \frac{2\beta\lambda_t+\alpha}{\lambda_b^2}$.$\beta$ and $\alpha$ can be determined based on $I_1 = 2\lambda_b^2\lambda_t$ and $\lambda_3 = \lambda_t$ separately. This give $s$ versus $\lambda_b$ curves for equal-biaxial loading as shown in Figure 11.

### Methods

#### E Tensile Tests

##### Custom micromechanical tensile stage

The custom tensile stage is shown as a 3D CAD representation in Figure 12. This stage includes the following components: (1) Steel bottom plate; (2) Newport MS-125-XYZ miniature linear stage; (3) Aluminum connection plate1; (4) Newport AG-LS25 piezo motor driven linear stage; (5) Aluminum connection plate 2; (6) Aluminum rod fixture; (7) L-shape fixture; (8) Futek LSB200 load cell; (9) Aluminum L-shape connection.

##### Leaf epidermal peel preparation

A mixture of 5g gelatin (Fisher, G8500) and 10g water was heated until the gelatin dissolved completely and then maintained at 80°C. The mixture was applied to a piece of cloth, and a dissected leaf was placed quickly onto the gelatin-applied area (Figure 5A). The cloth was folded and gently pressed to ensure the gelatin mixture made full contact with both surfaces of the leaf. After a half-hour curing process at room temperature, the folded cloth was unfolded by quickly pulling apart the two ends, separating the abaxial epidermal peel from the leaf (Figure 5B). A soft wet brush was used to gently sweep the peeled surface to remove any mesophyll cells attached to the abaxial epidermal peel. The cloth was then immersed in water for 20 seconds to reduce adhesion between the leaf and gelatin. Finally, the leaf epidermal peel was isolated from the cloth using tweezers.

##### Fluorescent bead dilution

We diluted the fluorescent bead suspension with Milli-Q Water by factors of 10, 100, and 500. The different concentrations did not have a noticeable influence on the mechanical behavior of the epidermal sample. A dilution factor of 100 provided the best pattern for DIC analysis.

#### F Scanning Electron Microscopy (SEM) imaging

Leaf epidermal peels were obtained as described in the tensile test methods and mounted to coverglass with the inner surface facing up. The peels were fixed in FAA (50% ethanol, 3.7% formaldehyde, 5% acetic acid) for 4 hours at room temperature. Following fixation, the peels were subjected to a series of increasing ethanol concentrations (50%, 50%, 60%, 70%, 80%, 90%, 95%, 100%, 100%, 100%) for 30 minutes each at room temperature and kept in 100% ethanol overnight at 4°C. On the second day, the peels were critical point dried and sputter-coated with gold palladium for 2 minutes to a thickness of approximately 40 nm. SEM imaging was performed using a Zeiss Gemini 500 Scanning Electron Microscope with EHT = 1.00 kV.

#### G Finite element simulations

##### Extracting the cellular structure from the confocal images

First, the maximum intensity Z projection image was obtained from the Zeiss z-stack image. This projection image was then segmented using Watershed segmentation from the MorphoLibJ plugin in ImageJ software. The outline of the cells was traced (Centerline tracing) from the segmented image, and any gaps were fixed using the Snap Object Points plugin and manual adjustments. The resulting cellular structure was exported as a .dxf file in Inkscape, which was then imported into ABAQUS Standard(2018). In ABAQUS, the cellular structure was constructed by extruding the cell outline to form the side walls and adding flat top and bottom walls.

##### Finite element type

The constructed cellular structure was meshed with 4-node reduced integration shell elements (S4R with hourglass control and finite membrane strains) with thickness of cell walls corresponding to thickness of the shell. The seed distance was set to 4 µm for the top/bottom walls and 2 µm for the side walls, based on convergence studies.

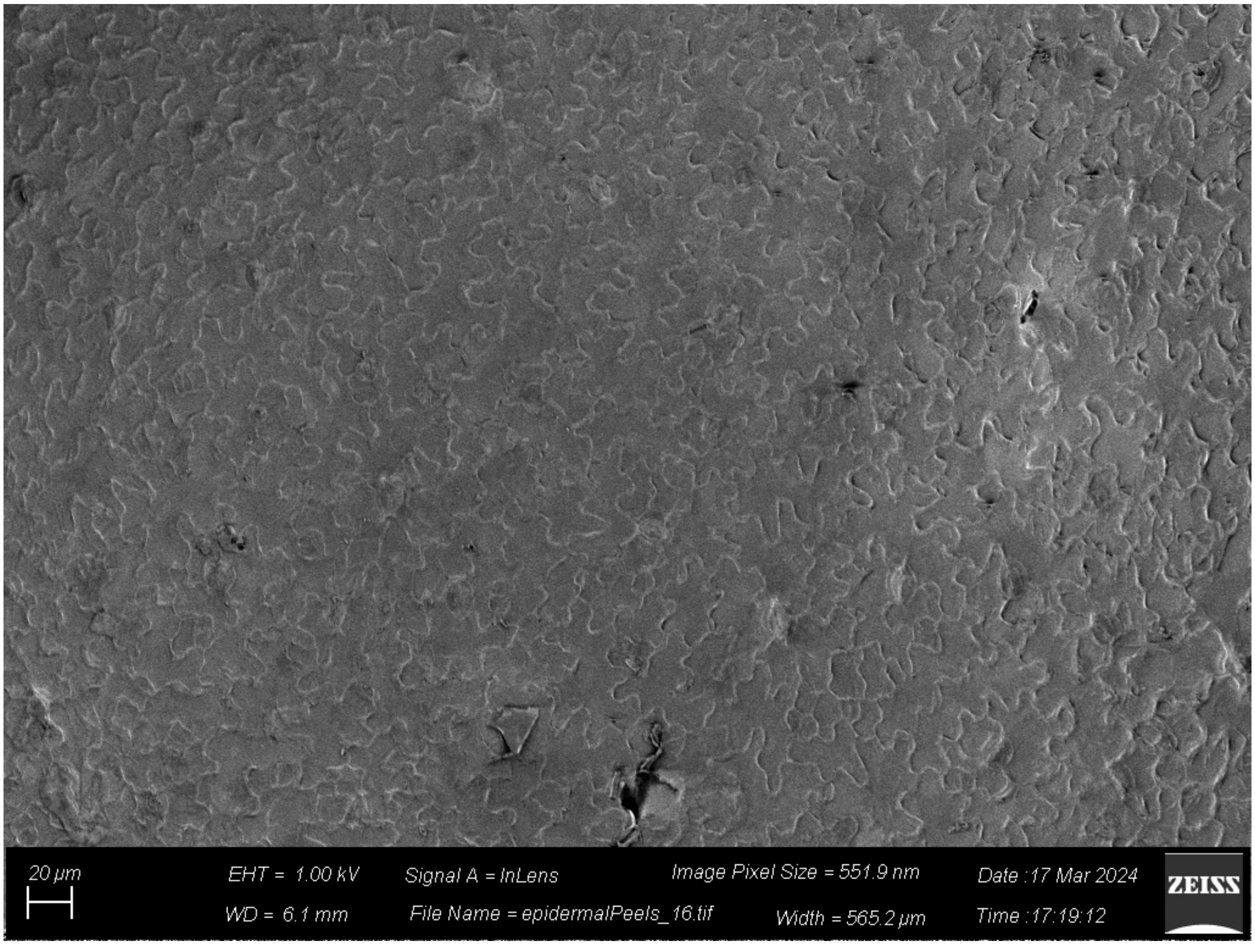


Figure 1: SEM image of leaf epidermal peel with the inner surface facing upwards.

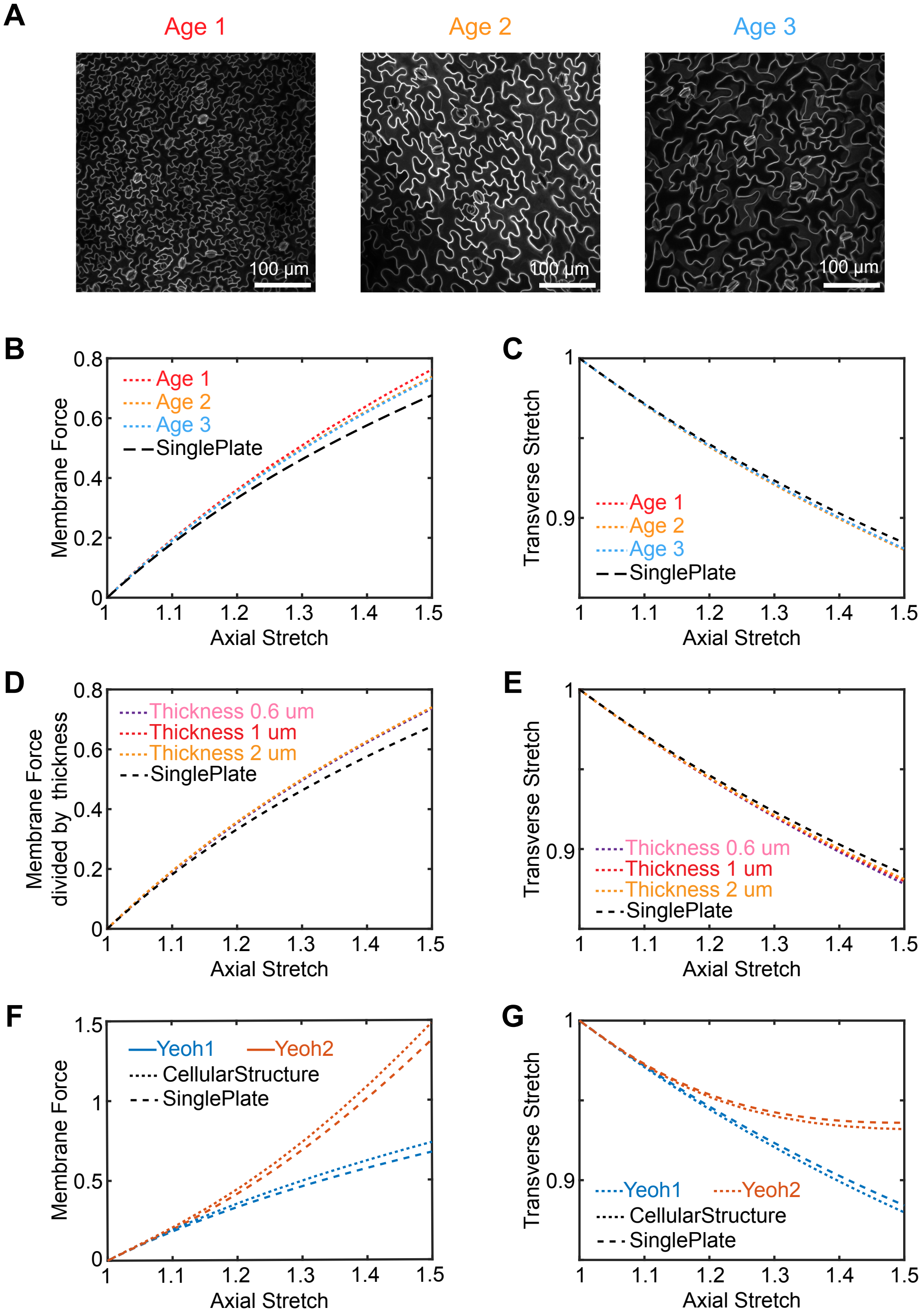


Figure 2: Cellular structure has little impact on overall mechanical behavior in cases of varying the cell size of the cellular structure, the thicknesses of the cell wall, and the material models of the cell wall. (A) Confocal images of abaxial epidermal cells from leaves at various developmental stages showing different cell sizes: small (12d), medium (18d), and large (25d) cell sizes; (B) Membrane force versus axial stretch curves and (C)Transverse stretch versus axial stretch curves of FEM simulation of three cellular structures constructed from three configuration and a single plate; (D) Membrane force versus axial stretch curves and (E)Transverse stretch versus axial stretch curves of FEM simulation of cellular structures with three different thicknesses and a single plate; (F) Membrane force versus axial stretch curves and (G)Transverse stretch versus axial stretch curves of FEM simulation of a cellular structure and a single plate using the Yeoh hyperelastic model with two different material parameters.

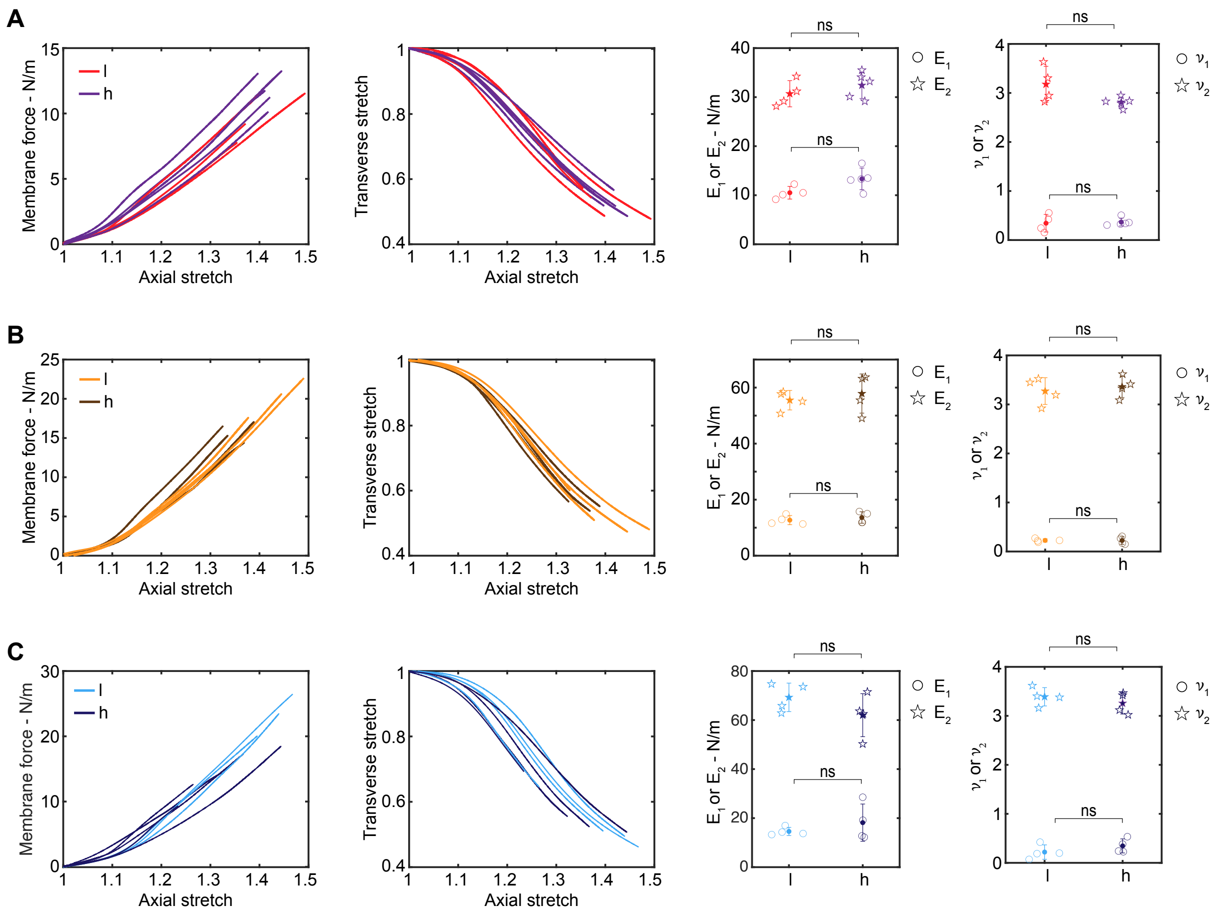


Figure 3: Comparison of monotonic tensile test results for wild type in proximal-distal (l) and medial-lateral (h) directions at three developmental stages: (A) 12 days, (B) 18 days, and (C) 25 days. This includes membrane force versus axial stretch curves, transverse stretch versus axial stretch curves, initial stiffness ($E_1$), final stiffness ($E_2$), initial Poisson's ratio ($\nu_1$), and peak Poisson's ratio ($\nu_2$), compared between directions for each stage respectively.

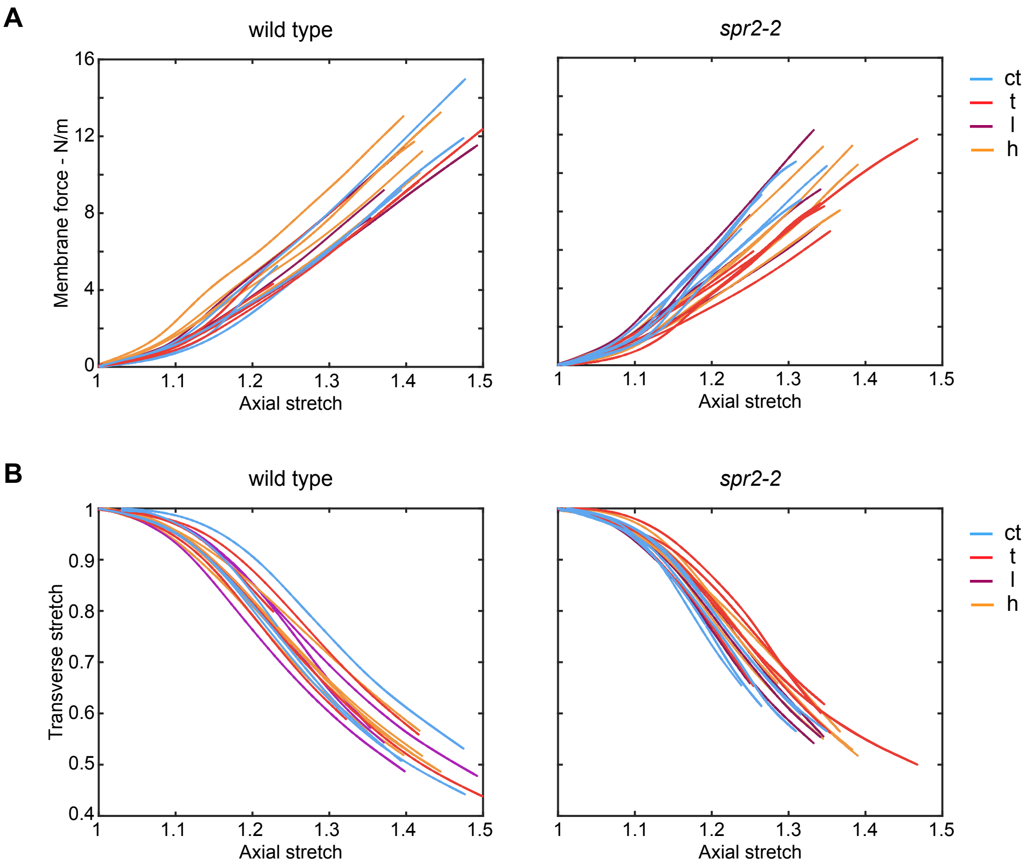


Figure 4: Comparison of monotonic tensile test results for wild type and *spr2-2* of 12d in four orientations: counterclockwise tilted 45 degrees relative to the midrib (ct), clockwise tilted 45 degrees relative to the midrib (t), proximal-distal (l), and medial-lateral (h): (A) membrane force versus axial stretch curves and (B) transverse stretch versus axial stretch curves.

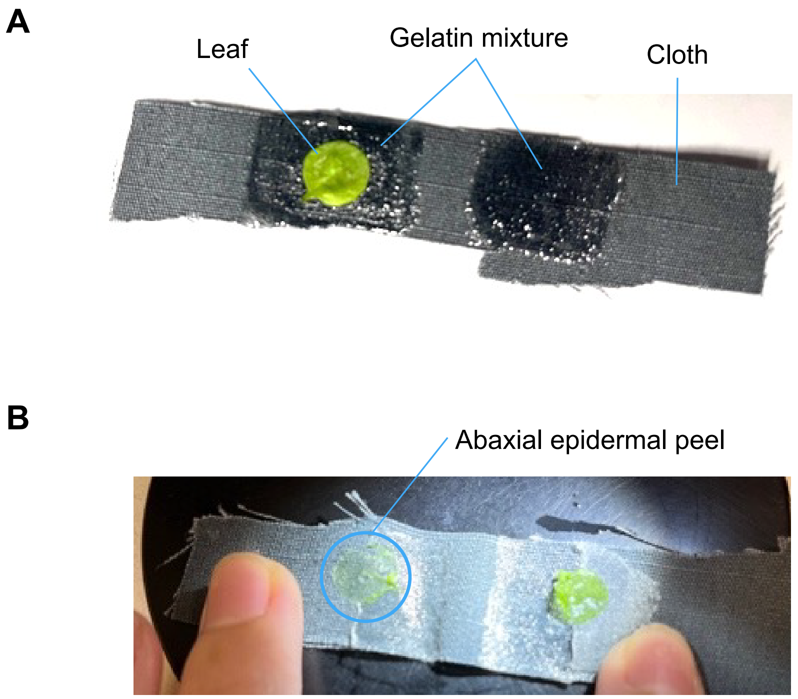


Figure 5: Obtaining leaf epidermal peels using gelatin-coated cloth. (A) The cloth was coated with gelatin mixture, and the leaf was placed on the gelatin-coated area. (B) After peeling, the abaxial epidermal peel was obtained.

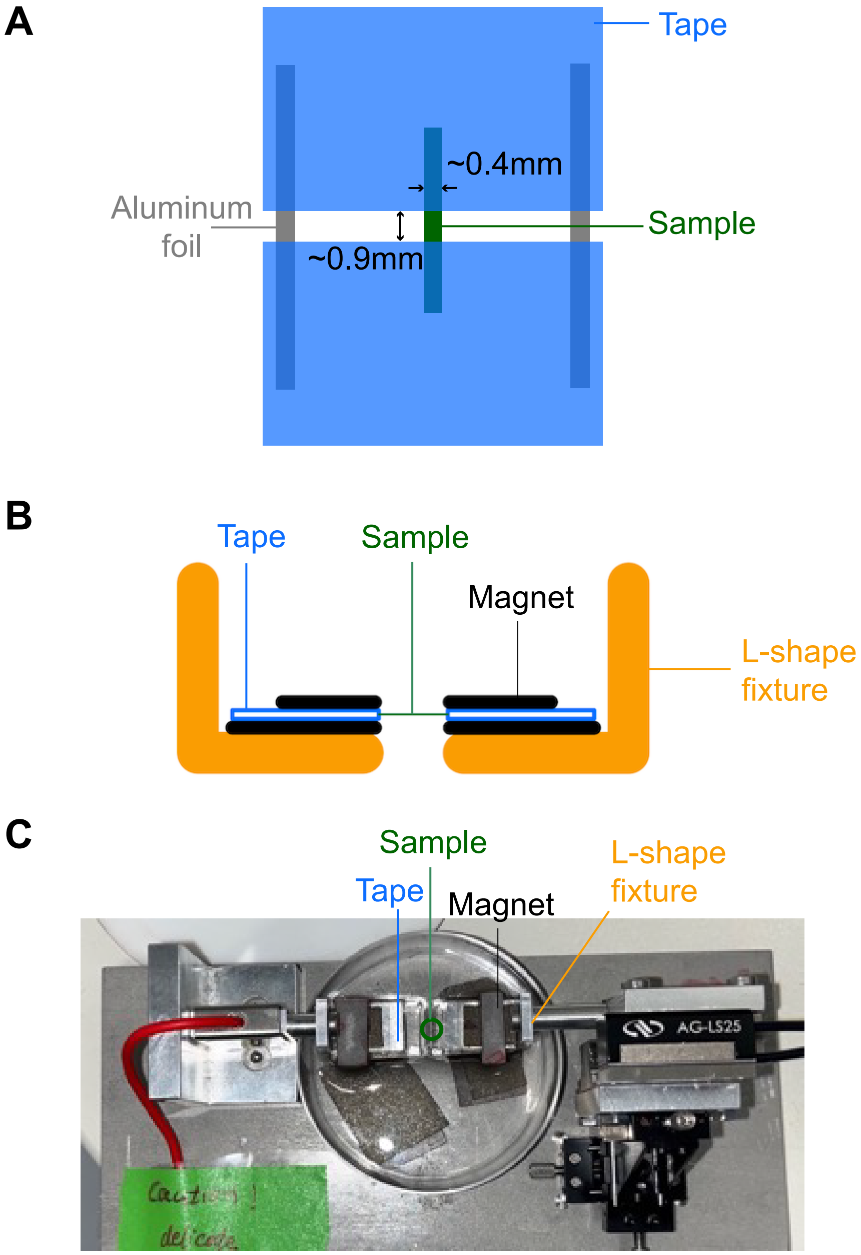


Figure 6: Attaching the sample on the micro-mechanical tensile stage. (A) Gripping a sample using tapes. (B) Schematic of using magnets to attach the tape-gripped sample to the L-shape fixture of the stage. (C) Top view of the tensile stage with the sample attached.

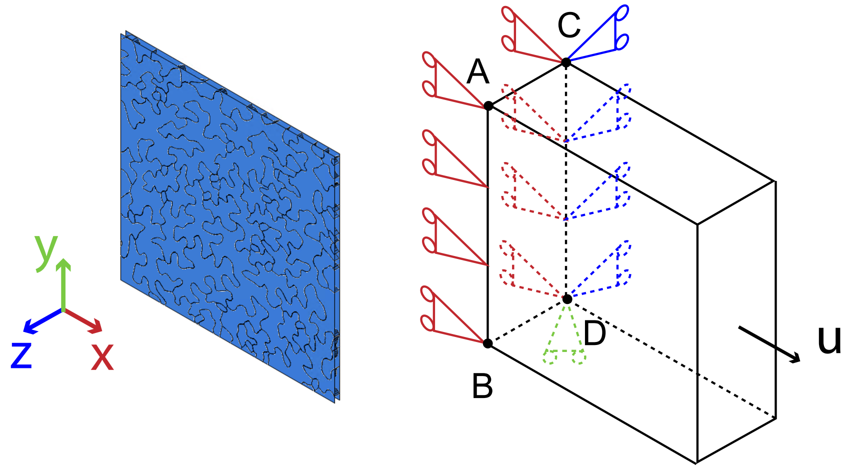


Figure 7: Boundary conditions for FEM simulations. $x = 0$ on boundary AB, $x = 0$ and $y = 0$ on boundary CD, and point D was fixed, and incremental displacement values $u$ were applied to the opposite surface.

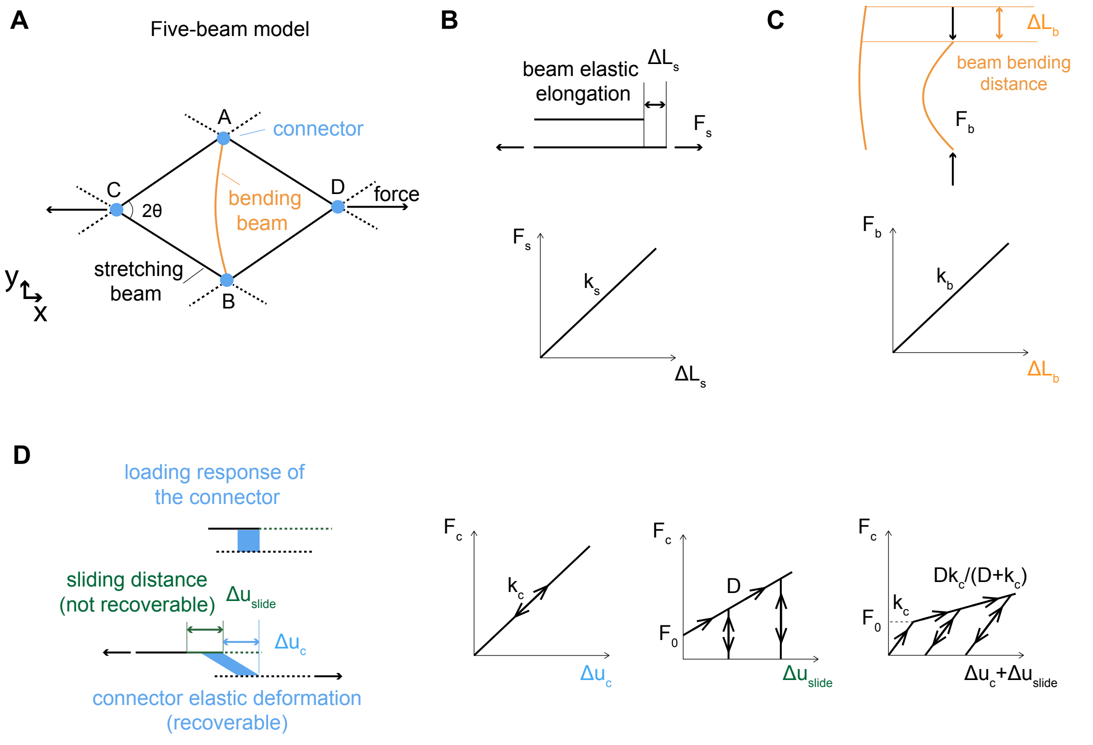


Figure 8: Five-beam model and the force-displacement responses for each components. (A) schematic of the five model (B) stretching beams (C) bending beam (D) connectors of the five-beam model

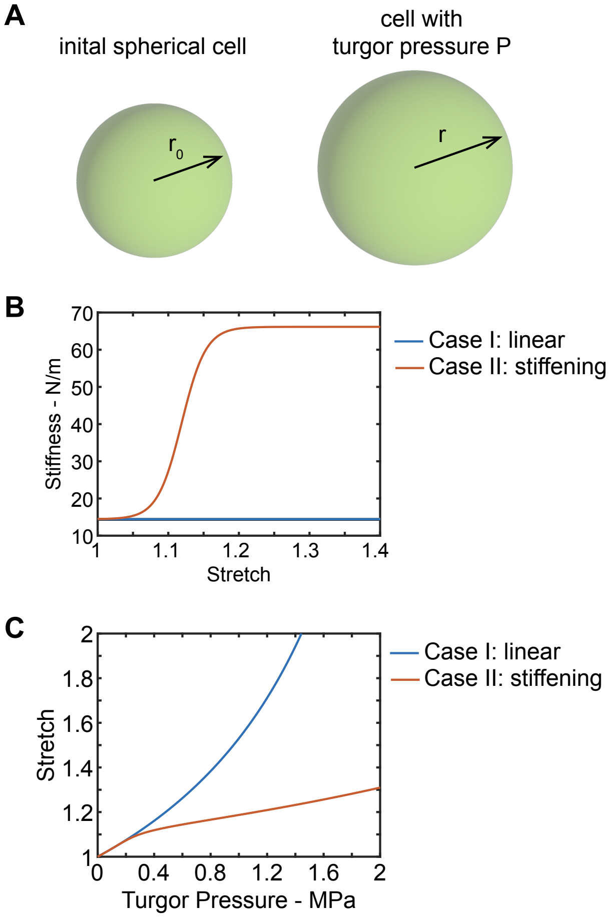


Figure 9: The deformation of spherical cell under turgor pressure for linear behavior and nonlinear stiffening behavior of the cell wall. (A) initial spherical cell and spherical cell with turgor pressure (B) the stiffness versus the stretch curves (C) stretch of the cell wall versus turgor pressure for linear behavior and nonlinear stiffening behavior.

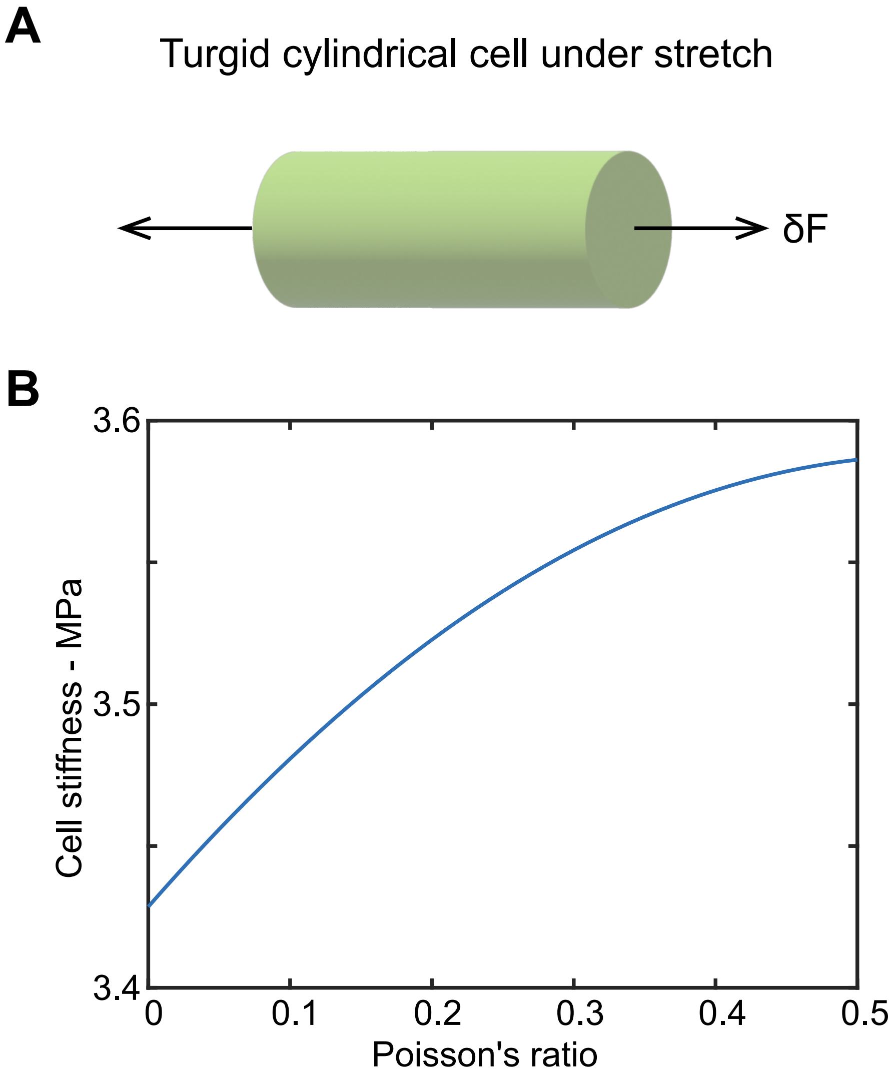


Figure 10: Poisson's ratio influences the stiffness of a turgid cylindrical cell. (A) A turgid cylindrical cell under axial loading. (B) Stiffness of the cell versus Poisson's ratio curve.

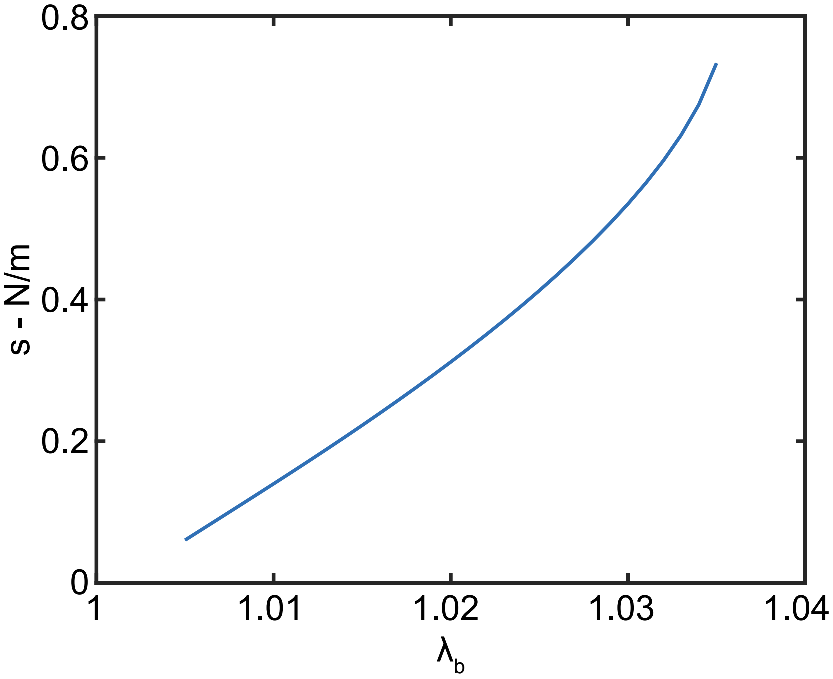


Figure 11: Membrane force versus stretch under equal-biaxial loading.

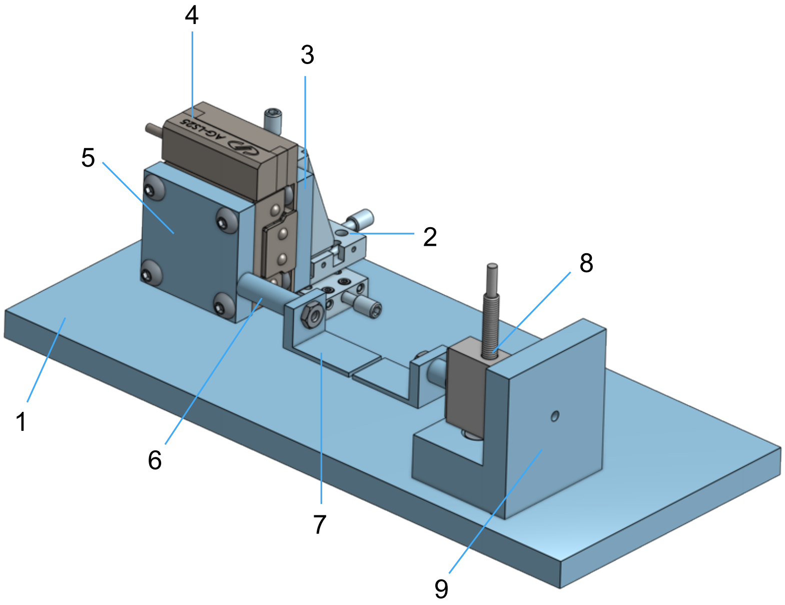


Figure 12: 3D CAD drawing of the micro-mechanical tensile stages: (1) Steel bottom plate; (2) Newport MS-125-XYZ miniature linear stage; (3) Aluminum connection plate1; (4) Newport AG-LS25 piezo motor driven linear stage; (5) Aluminum connection plate 2; (6) Aluminum rod fixture; (7) L-shape fixture; (8) Futek LSB200 load cell; (9) Aluminum L-shape connection.

Table 1: Parameters of material models used in FEM simulations.

| Models | C10 | C20 | C30 | D1 | D2 | D3 |
|---|---|---|---|---|---|---|
| neo-Hookean | 0.1923 | NA | NA | 2.4 | NA | NA |
| Yeoh 1 | 0.1923 | -0.002 | 0.002 | 2.4 | 0 | 0 |
| Yeoh 2 | 0.1923 | 0.3846 | 0 | 2.4 | 0 | 0 |

#### Movie captions

**Movie 1**: Digital image correlation tracking on the stretching sample. The sample was coated with green fluorescent beads, and the heat map shows the percentage of deformation in the stretch direction.
**Movie 2**: Finite element simulation of stretching single plate. The heat map was showing the stress level in the stretching direction.
**Movie 3**: Finite element simulation of cellular structure. The heat map was showing the stress level in the stretching direction.
